# Notes on Synthesis of perdeutero-5-^13^C,5,5,5-trifluoroisoleucine VI^56^

**DOI:** 10.1101/140681

**Authors:** D.G. Naugler, R. S. Prosser

## Abstract

The ^13^CF_3_ group is a promising label for heteronuclear (^19^F,^13^C) NMR studies of proteins. Desirable locations for this NMR spin label include the branched chain amino acid methyl groups. It is known that replacement of CH_3_ by CF_3_ at such locations preserves protein structure and function and enhances stability. ^13^CF_3_ may be introduced at the a position of isoleucine and incorporated biosynthetically in highly deuterated proteins. This paper reports our work in synthesis and purification of 5,5,5-trifluoroisoleucine, its perdeutero and 5-^13^C versions and of 2-^13^C-trifluoroacetate and its utility as a precursor for introduction of the ^13^CF_3_ group into proteins.

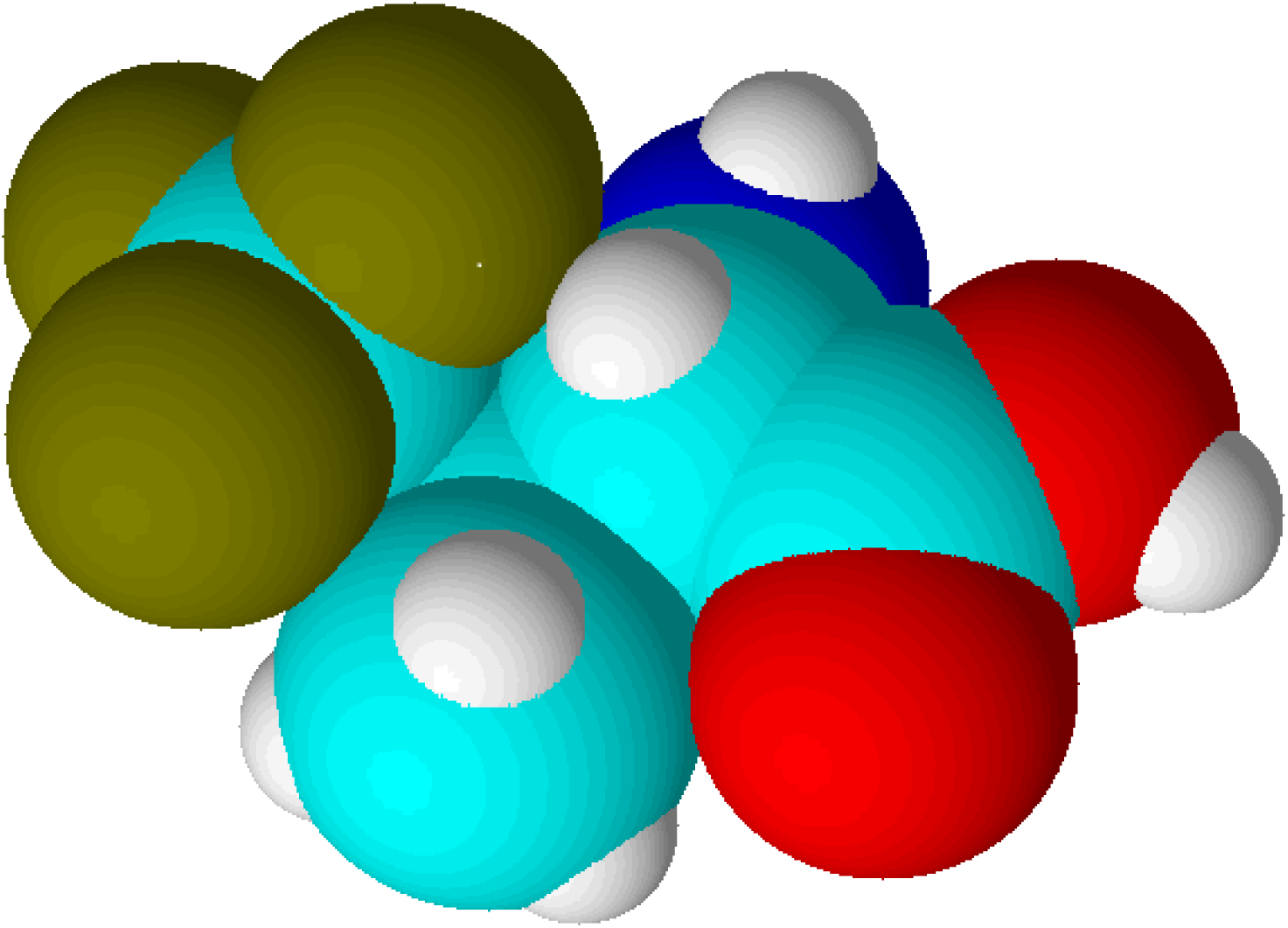

## Introduction

### Fluorine NMR and labeling strategies in proteins

Fluorine NMR spectroscopy is a powerful method for the study of both structure and dynamics of proteins and their interactions with other proteins or ligands^1,2,3,4^. Because of the ability of the ^19^F lone-pair electrons to participate in non-bonded interactions with the local environment, ^19^F chemical shifts are sensitive to changes in van der Waals contacts, electrostatic fields and hydrogen bonding. As such, ^19^F chemical shifts (or changes in shifts) are often indicative of conformational changes^5,6,7,8^, binding^9^, and protein folding or unfolding events^10,11,12^. Fluorinated probes are also frequently used to assess solvent exposure, via chemical shift changes or relaxation effects resulting from: 1) substituting H_2_O for ^2^H_2_O^2,14^ paramagnetic additives such as Gd^3+^:EDTA^2^ heteronuclear nuclear Overhauser effects^13^. In membranous systems, analogous paramagnetic effects are also observed upon addition of nitroxide spin-labels^14^ or dissolved oxygen^15,16^, facilitating the study of topology and immersion depth via fluorinated probes. Finally, associated ^19^F spin-spin and spin-lattice relaxation rates are useful for studying conformational dynamics over a wide range of timescales, due to the significant chemical shift dispersion, shift anisotropy, and large heteronuclear dipolar relaxation terms^17^, ^18^, ^19^, ^20^.

Fluorine labeling of proteins is achieved in many ways. Via biosynthetic means, monofluorinated versions of tyrosine, phenylalanine, and tryptophan may be substituted for their nonfluorinated equivalents, often with little effect on overall expression yields^2,4^. Fluorinated versions of methionine^7,21^, proline^22^, leucine^23^, and isoleucine^24^ have also been successfully incorporated into proteins. An alternative approach to ^19^F labeling of proteins is to make use of a thiol specific fluorinated probe, which frequently consists of a terminal trifluoromethyl group^4^. In this way, a fluorine tag may be placed at virtually any site in the protein, using successive single cysteine mutations of the protein under study. Thus, the majority of ^19^F labels used in protein NMR fall under the category of isotopic fluoroaromatic or trifluoromethyl species. For very large proteins, spectral overlap may become problematic for biosynthetically labeled proteins. However, a doubly (^13^C, ^19^F) labeled amino acid should provide greater resolution in two-dimensional (^19^F,^13^C) NMR spectra, with further possibilities of assignment without mutational analysis. Moreover, such two-dimensional NMR schemes should benefit from the relatively large one bond (^13^C,^19^F) coupling which is between 265 and 285 Hz for both fluoroaromatics and trifluoromethyl groups. Finally, the possibilities for studying dynamics from a ^13^C,^19^F pair encompasses a much greater range, since various zero, single, and double quantum coherences in addition to Zeeman and two-spin longitudinal order, may be separately evolved and studied^25,26^. In this paper, we present a method for the preparation of perdeuterated isoleucine, in which the terminal trifluoromethyl group consists of a ^13^C-^19^F pair. The motivation for this work is to develop a useful doubly labeled species for subsequent nD NMR studies of proteins, whose isoleucine residues have been fluorinated.

### Advantages of a trifluoromethyl group

The trifluoromethyl group is expected to be a useful probe of molecular structure and dynamics, particularly in the hydrophobic core of proteins, at the interface between protein complexes, and in the membrane or detergent interior in studies of integral membrane proteins. Expressed within proteins, the CF_3_ group offers additional benefits of sensitivity and relatively long transverse relaxation times. However, the inherent slow rotational tumbling associated with large proteins or protein complexes, and membrane proteins, results in line broadening and reduced sensitivity. ^19^F spin labels also suffer extensively from dipolar relaxation with nearby proton spins of the protein^2^, which may be largely avoided by extensive deuteration. Furthermore, in situations where ^13^C,^19^F two-dimensional NMR schemes are employed, the use of transverse relaxation optimized spectroscopy (TROSY) techniques^27,28^ may be considered. The TROSY effect in methyl groups, results from interference between intra-methyl dipolar interactions^28^. As such, the effect is independent of field, to the extent that chemical shift anisotropy does not contribute to relaxation. Since the geometry of the trifluoromethyl group is like that of a CH_3_ group, while the gyromagnetic ratio of the ^19^F nucleus is 0.83 times that of ^1^H, the methyl TROSY effect would be expected to be preserved in appropriate (^19^F,^13^C) two-dimensional schemes. In particular, in the rigid limit, the maximum peak intensities in the (^1^H,^13^C) HMQC are predicted to depend on terms which are derived from relaxation via reorientation of either the CF or FF intramolecular bonds *i.e.*, transverse rates proportional to 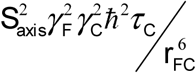 and, to a lesser extent,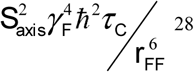 In this case, *S* _*axis*_ and τ_c_ represent the local order parameter of the methyl rotor and the global correlation time associated with rotational tumbling of the protein. Finally, γ_C_ and γ_F_ represent the magnetogyric ratios of ^13^C and ^19^F, respectively while **r**_CF_ and **r**_FF_ designate the intramolecular bond lengths. In the (^19^F,^13^C) HMQC, where we estimate the CF and FF intramolecular bond lengths in a trifluoromethyl group to be 1.33 Å and 2.12 Å respectively, the above transverse rates are predicted to be more than three times smaller than those for the CH_3_ groups, in the absence of external dipolar relaxation or relaxation due to chemical shift anisotropy.

### The trifluoromethyl probe in isoleucine

Considering these anticipated advantages for 1D and 2D NMR, we have developed a protocol for the synthesis and purification of perdeuterated 5,5,5-trifluoroisoleucine, in which the carbon nucleus of the trifluoromethyl group is ^13^C enriched. Incorporation, using a cell-free protein expression technique, is reported^55^. We also describe herein a synthesis strategy for 2-^13^C-trifluoroacetate and purification of the ammonium salt (or hypothetically CF_3_CO_2_H), to produce perdeutero 5-^13^C-5,5,5-trifluoroisoleucine. This report is intended to communicate some of the subtleties involved in these efforts.

Isoleucine has some additional features that make it attractive for (^19^F, ^13^C) double labeling. At neutral pH, isoleucine is the most hydrophobic of the aliphatic side chain amino acids and it is often highly conserved at positions involved in hydrophobic interactions. Consequently, isoleucine often functions at the hydrophobic binding cleft. For example, isoleucine residues 737 and 898 in the human androgen receptor ligand binding domain mediates interdomain communication with the NH_2_-terminal domain, which in turn mediates transcriptional activation^29^. Calmodulin provides another example. ^1^H NMR studies of amide proton exchange rates of Ca^2+^-saturated calmodulin and a Ca^2+^-saturated calmodulin-mastoparan complex showed a reduction in solvent accessibility of Ile27 upon mastoparan binding^30^. Genomic analysis of membrane protein families informs us of amino acid abundances and conserved motifs. The most abundant amino acids in transmembrane regions are leucine, isoleucine, valine, phenylalanine, alanine, glycine, serine, and threonine. Taken together, these amino acids account for 75% of the amino acids in transmembrane regions. In contrast, isoleucine (10% abundance versus 4% conserved), valine (8% versus 4%), methionine (4% versus 1%) and threonine (7% versus 4%) are less prevalent in conserved positions^31^. Large hydrophobic (Phe, Leu, Ile, Val) residues show a clear preference for the protein surfaces facing the lipids for β-barrels, but in α-helical proteins, no such preference is seen, with these residues equally distributed between the interior and the surface of the protein^32^.

### Synthesis outline

The scheme shown below shows the route used to make 5,5,5-trifluoroisoleucine as a racemic mixture of diastereomers^33^. This scheme is modified for perdeuteration by substitution with cyanoacetic acid-d_3_, acetone-d_6_, CF_3_CO_2_D, ammonium-d_4_ acetate-d_3_ and D_2_/Pd. Modification of this scheme for the ^13^CF_3_ amino acid requires electrochemical trifluoromethylation using ^13^CF_3_CO_2_^−^.

### Experimental

Melting points are uncorrected. ^1^H, ^19^F and ^13^C NMR spectra were recorded on Varian Gemini 200 MHz and Unity Inova 600 MHz spectrometers. ^2^H NMR spectra were recorded on the Varian Unity Inova 600 MHz spectrometer. UV spectra were recorded using a Biochrom Libra S22 UV/Visible spectrophotometer. IR spectra were recorded using a Nicolet Avatar 360 E.S.P. FT-IR spectrometer. GC/MS was done using a Shimadzu GCMS-QP5050 system. LC/MS was performed using a system comprised of a Waters/Alliance 2690 liquid chromatography module and a Waters Micromass ZQ ESI mass spectrometer. Hydrogenation was performed with a Parr high-pressure hydrogenation apparatus. Electrochemical trifluoromethylation was performed with a Sargent Slomin analyzer with (one spinning) platinum mesh electrodes.

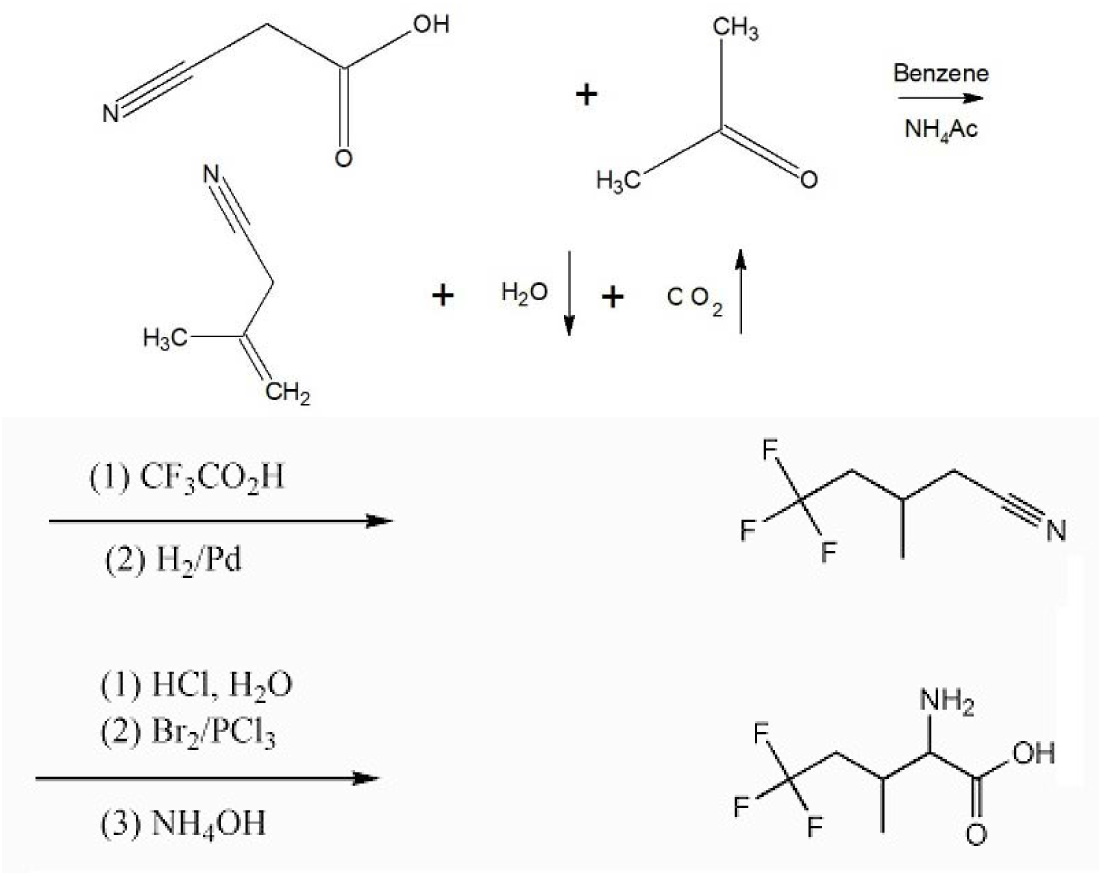

### Synthesis of methallylcyanide and methallylcyanide-d_7_

Synthesis of methallylcyanide and methallylcyanide-d_7_ relies on eactive hydrogen chemistry, *i.e.* exchangeable protons/deuterons. Wang et al.^27^ cite a synthesis reported by Marson *et al.*^34^. Cyanoacetic acid (e.g. 50.0 g, 0.59 mol), acetone (34.2 g, 0.59 mol), and ammonium acetate (4.0 g, 0.05 mol) (or their perdeutero counterparts) in dry benzene (100 mL) were refluxed with a Dean-Stark trap. Reflux was performed until one equivalent of water (or D_2_O) was collected in the trap. Due to the cost of perdeuteroammonium acetate, a minimal amount of this catalyst was used. The quantity was observed not to be critical to yield. One gram of ammonium-d_4_ acetate-d_3_ will suffice for the synthesis of methallylcyanide-d_7_. The Dean-Stark unit was replaced with a distillation head and the fraction collected between 110 °C and 115 °C.

Compton et al.^35^ reports the existence of both 3-methyl-2-butenenitrile and 3-methyl-3-butenenitrile in a sample of methallylcyanide. This is consistent with an equilibrium arising from eactive hydrogen chemistry as evidenced by the ^2^H NMR spectrum in Figure 1, below. The predicted boiling points for these isomers are within 2°C, so they cannot be separated by fractional distillation. Accordingly, this product is properly called methallylcyanide^33^ rather than 3-methyl-but-3-enenitrile^24^. Appreciation of the active hydrogen nature of this product is required to make methallylcyanide-d_7_.

**Figure 1.**
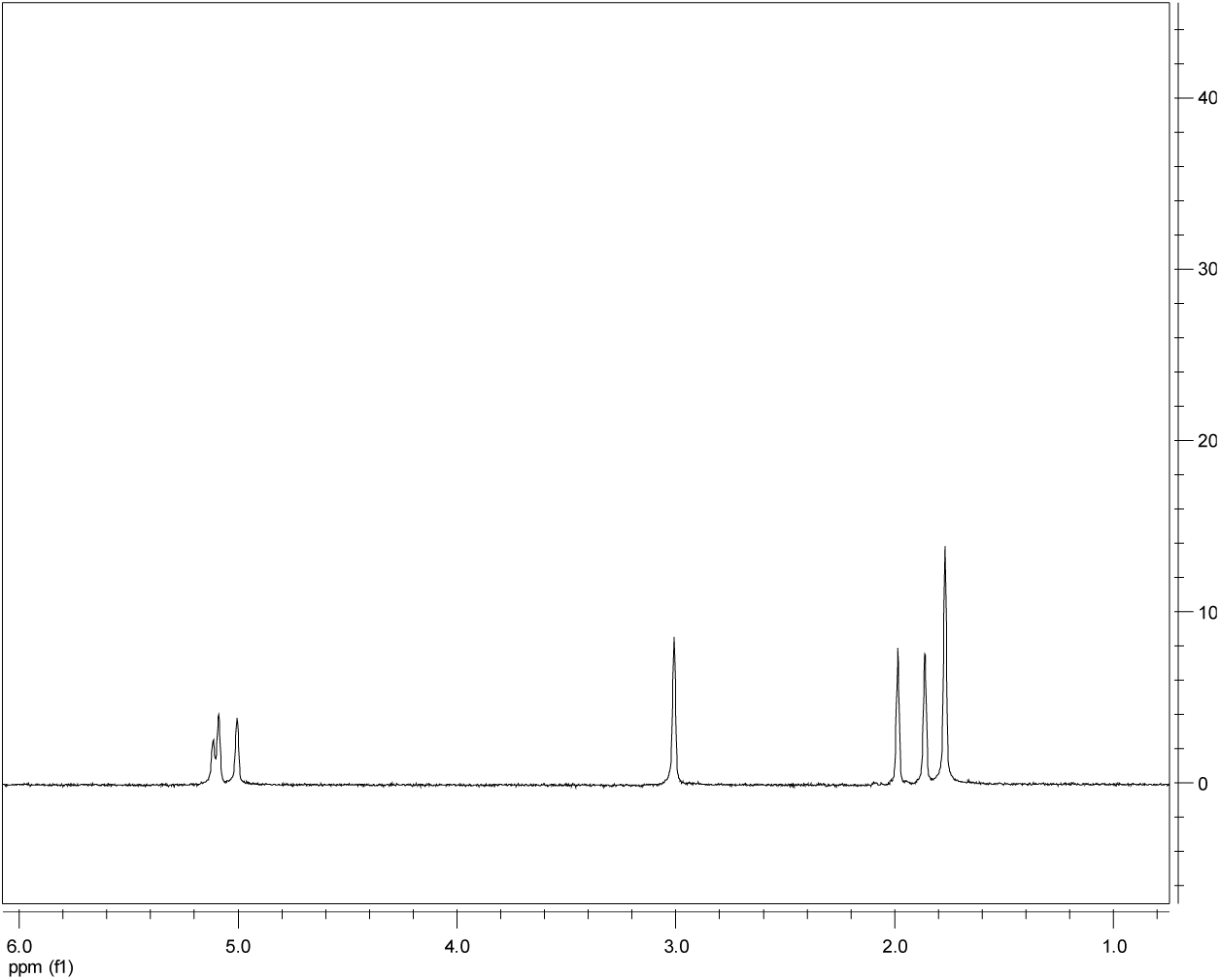
Deuterium NMR spectrum of methyallylcyanide-d_7_, an equilibrium mixture of 3-methyl-2-butenenitrile-d_7_ and 3-methyl-3-butenenitrile-d_7_. The predicted boiling points of these two components are within 2 °C, thus these isomers cannot be separated by fractional distillation.

**Figure 2.**
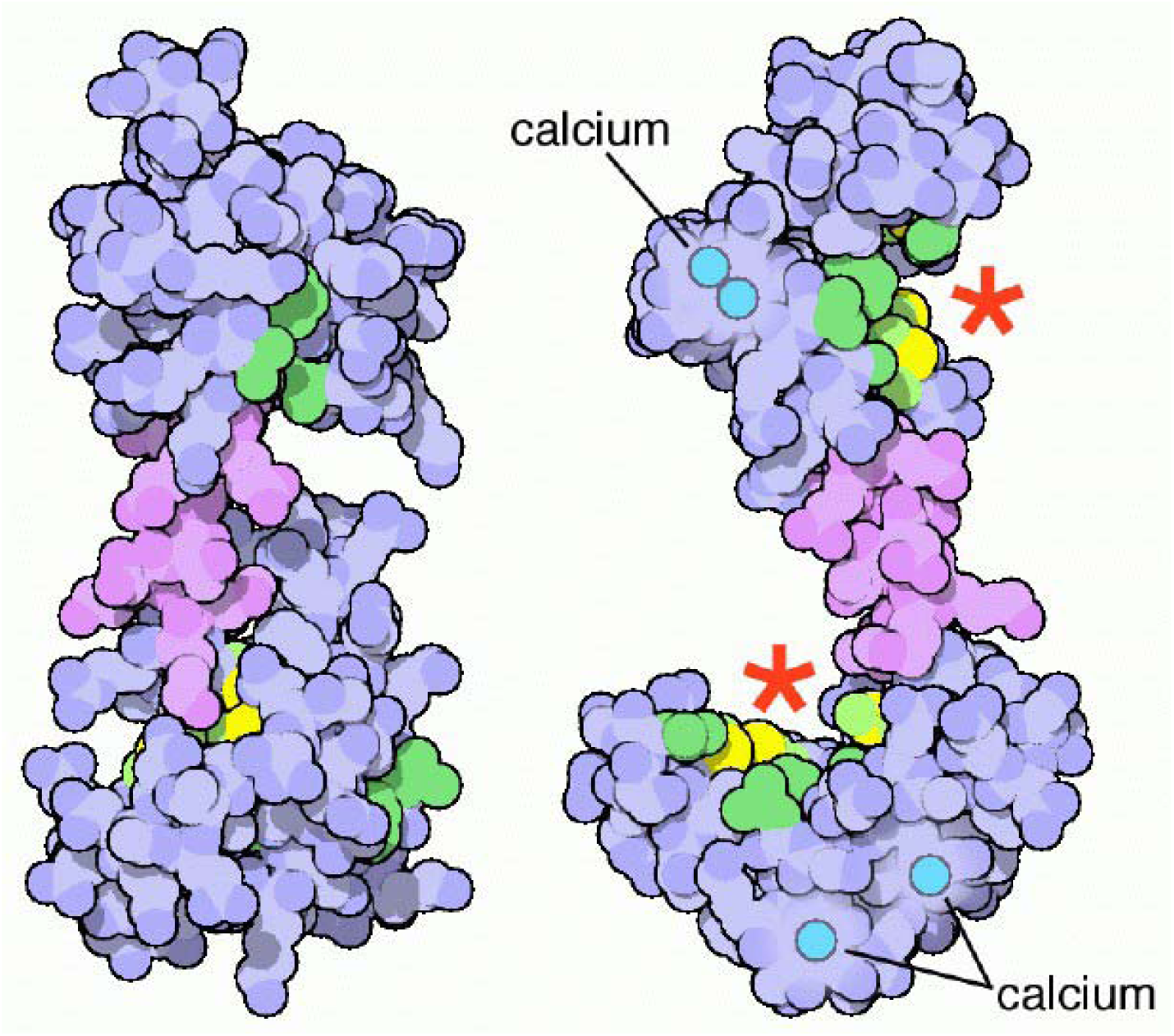
These images show conformational changes in calmodulin. On the left is calmodulin without calcium and on the right, is calmodulin with calcium. Sites that bind target proteins are indicated by red stars. These images are from the RCSB Protein Data Bank. <u>http://pdb101.rcsb.org/motm/44</u>, <u>https://meshb.nlm.nih.gov/record/ui?name=Calmodulin</u>

Synthesis of methallylcyanide-d_7_ requires the preparation of cyanoacetic acid-d_3_, first and then its condensation with acetone-d_6_, catalyzed by ammonium-d_4_ acetate-d_3_. 50 grams of cyanoacetic acid was dissolved with warming in a minimum quantity of D_2_O, then left overnight to affect exchange. Excess H_2_O/D_2_O was subsequently removed with a rotary-evaporator, and the wet crystals further dried overnight in a vacuum desiccator. Because not all H_2_O/D_2_O can be removed by this procedure, subsequent steps require less D_2_O to affect dissolution. The proton/deuteron exchange procedure was executed three times and the product submitted for mass spectral analysis. Three more H/D exchange steps were performed. The resulting cyanoacetic acid-d_3_ was condensed with a molar excess of actetone-d_6_ in the presence of ammonium-d_4_ acetate-d_3_ and the water of condensation was analyzed by ^1^H NMR. The trace aqueous solubility of benzene is known^36^. Comparison of integrated peak intensities showed that the water of condensation contained 0.6% H_2_O and 99.4% D_2_O. Given that all H/D sites are exchangeable, it is concluded that methallylcyanide-d_7_ obtained after decarboxylation and distillation of the condensation product was 99.4% isotopically pure. This isotopic purity is consistent with that of the reagents used. See the ^2^H NMR spectrum in Figure 1. below.

### Electrochemical trifluoromethylation of methallylcyanide-d_7_

Muller^33^ chose to perform electrochemical trifluoromethylation of methallylcyanide in aqueous 90% methanol between platinum electrodes whereas we chose a later system^37^ of CH_3_CN and H_2_O (8:1). An electrochemical solvent system of acetonitrile and water necessitates further use of acetonitrile and D_2_O for methallylcyanide-d_7_.

Because of hydrogen exchange chemistry of methallylcyanide-d_7_, special care needs to be applied to the adaptation of this reaction to a perdeuterated substrate. H_2_O needs to be replaced by D_2_O. Both MeOD and CH_3_CN methyls ought to be resistant to H/D exchange under the conditions of the reaction but CH_3_CN is cheaper than MeOD and so is a better choice. Muller^33^ chose to use a 73% molar excess of CF_3_CO_2_H relative to methallylcyanide and to use a slight coulombic excess. Dmowski^37^ chose to use a 20 times molar excess of TFA relative to substrate and 1.5 Faradays per mole of TFA. Because both methallylcyanide-d_7_ and 2-^13^C-trifluoracetate are very expensive these should be used in 1:1 molar ratio and with 50% excess current. Both authors achieved mild basic condition with 10% Na.

We used a Sargent Slomin S-29460 electrolytic analyzer, for radical trifluoromethylation. Both rotating and stationary electrodes were platinum. The rotating platinum electrode was chosen to be the anode. At the anode, reactions CF_3_CO_2_^−^ → CF3CO_2_. + e^−^ → CF_3_· + CO_2_↑ and at the cathode 2H^+^ + 2e^−^ → H_2_↑, both yield gases. The spinning electrode helps stirring and shakes off gas bubbles. Upon entering the limiting current region increasing the applied voltage is counterproductive. Gas production observation aids adjustment. The S-29460 electrolytic analyzer displays voltage and amps. Muller^33^ recommends a stoichiometric excess of coulombs to drive the reaction to completion. The electro-synthesis was performed in an open beaker, using an ice bath for cooling. This was conducted in a fume hood.

Muller s workup^33^ was followed, as described below. For methallylcyanide-d_7_, H_2_O should be replaced by D_2_O, H_2_ by D_2_ and MeOH by MeOD or some other exchange resistant solvent such as CH_3_CN. An iron (steel) cathode would minimize loss of valuable substrate by electrolytic hydrogenation.

The mixture was poured into 700 mL of water, the dense oil collected, and the aqueous layer extracted with two 40 mL portions of dichloromethane. The combined organic layers from three identical runs were distilled to remove the solvent and then steam distilled. The non-aqueous layer was a mixture of 3-methyl-5,5,5-trifluoropentanonitrile, several isomeric 3-methyl-5,5,5-trifluoropentenonitriles, methallyl cyanide, and unidentified by-products. It was diluted with methanol and hydrogenated at low pressure over 5% Pd/C. Distillation afforded 66 g of nearly pure 3-methyl-5,5,5-trifluoropentanonitrile, b.p. 166-171 °C.

For the perdeutero route, once the various isomeric perdeutero-3-methyl-5,5,5-trifluoropentenonitriles are reduced by D_2_, exchange is no longer possible. Subsequent steps closely follow the protocol outlined by Muller as does this exposition. Quantities were adjusted proportionately.

The distillation product, 3-methyl-5,5,5-trifluoropentanonitrile was stirred with sufficient concentrated aqueous hydrochloric acid to bring most of the organic material into solution, for several days, then diluted with water and refluxed overnight. The separated organic layer was isolated, dried over Na_2_SO_4_, and distilled at 6 torr, giving 58.3 g of 3-methyl-5,5,5-trifluoropentanoic acid, b.p. 76-81 °C.

58.3 g (0.343 mol.) of 3-methyl-5,5,5-trifluoropentanoic acid, 19.1 mL of bromine and 0.6 mL of phosphorus trichloride were refluxed with a trap to absorb gaseous hydrogen bromide until the color of bromine had disappeared. On distilling at 6 torr, about 6 g of 3-methyl-5,5,5-trifluoropentanoic acid were recovered. 60 g of distillate boiling at 100-106 °C consisted mainly of nearly equal amounts of the two diastereomers of 2-bromo-3-methyl-5,5,5-trifluoropentanoic acid. This mixture was used for the preparation of 2-amino-3-methyl-5,5,5-trifluoropentanoic acid without further purification. 64.4 g (0.259 mol.) of this nearly pure 2-bromo-3-methyl-5,5,5-trifluoropentanoic acid and 225 mL of concentrated aqueous ammonia were cautiously mixed and stored in a closed flask at 44-48 °C for 5 days. The stopper was removed and the mixture gently heated with a water bath to drive off excess ammonia and reduce the volume to about 90 mL.

We observed the formation of polymeric materials and so we could not crystallize the product from our reaction mixture. Accordingly, we developed several purification methods discussed in the next section.

### Purification of a reaction mixture containing 2-amino-3-methyl-5,5,5-trifluoropentanoic acid

Two factors are likely to alter the physical properties of 5,5,5-trifluoroisoleucine relative to native isoleucine. The inductive effect of the CF_3_ group^38^ will make the amino acid and amino groups more acidic and the greater hydrophobicity of the CF_3_ group will enhance the hydrolytic stability of its polymers^39^.

In our hands, reaction of a mixture of diastereomers of 2-bromo-3-methyl-5,5,5-trifluoropentanoic acid with aqueous ammonia produced a product mixture that contained a significant quantity of polymeric material. Some of this material was readily soluble in diethyl ether and CDCl_3_. That fraction that was soluble in organic solvent could be hydrolyzed by dissolution in TFA followed by gradual addition of water. We found that the reaction mixture could be stabilized by formation of the TFA salt.

We first chose a method of chemical purification that was appropriate for the partial fluorous character of the amino acid^40,41,42^. The use of a C_8_F_17_ BOC derivative^42^ proved to be problematic because the only rational method for its hydrolysis was the use of TFA and we already had made the TFA salt. Exploitation of the C_8_F_17_ Cbz derivative^42^ proved to be more fruitful. A product mixture was dissolved in THF and a minimal quantity of water and reacted with a small molar excess of N-[4-(1H,1H,2H,2H-perfluorodecyl)benzyloxycarbonyloxy]succinimide. The resulting C_8_F_17_ Cbz derivative was purified by solid phase extraction on fluorous silica^40^. This derivative was subjected to atmospheric pressure hydrogenolysis over 5%Pd/C and the subsequent product mixture subjected to fluorous SPE. Lyophilization yielded a soluble product that was white. Incorporation of this product in calmodulin (Takeda & Kainosho, 2012)^55^ (M. Kainosho: private communication) using a cell free protein expression system proved that the product mixture contained ≤25% 5,5,5-trifluoro-L-isoleucine and proved the efficacy of the chemical purification.

During lyophilization, much of the product was lost due to sublimation. Sublimation has recently been revisited as a means for purification of amino acids^43^. We found that our chemically purified product mixture could be sublimed at 150 °C and 6 mm Hg, but that fractional sublimation would require better vacuum and temperature control.

Finally, we exploited ion exchange chromatography using cellulose phosphate to separate monomeric and polymeric fractions. Elution with distilled H_2_O yielded a microcrystalline fraction while elution with dilute aqueous ammonia yielded a waxy fraction identifiable as polymeric material. Refinement of ion exchange column purification should be developed using ion exchange TLC on cellulose phosphate paper.

We explored the use of analytical HPLC-MS first using an acetonitrile gradient in 0.1% aqueous TFA on a C_18_ column. 2-amino-3-methyl-5,5,5-trifluoropentanoic acid has a molecular weight of 185. We identified two separated peaks corresponding to [M+] and [M+H+] on an ESI-MS instrument. We conclude that these are the diastereomers and that differential inductive effects due to CF_3_ cause differing acid/base properties of the diastereomers. These peaks were followed at longer time by peaks due to polymers. We wished to explore the possibility of preparative scale chromatography where TFA would be counterproductive. An isocratic method was developed using 5% acetonitrile in 0.2% aqueous formic acid on an analytical C_18_ column.

### Ammonium 2-^13^C-trifluoroacetate

To embed the ^13^CF_3_ group into a synthetic scheme for 5,5,5-trifluoroisoleucine following the synthetic scheme above, a route to 2-^13^C-trifluoroacetate is required. Trifluoroacetic acid has been made via electrochemical fluorination^44^. Since this electrolysis would entail use of anhydrous hydrogen fluoride within a customized Teflon reaction vessel, a refrigeration unit, a high current power supply and a process control system, we chose to explore an alternate route involving halogen exchange with 2-^13^C-tribromoacetic acid. We devised a new route to 2-^13^C-tribromoacetic acid starting with 2-^13^C-ethanol, discussed below.

### Halogen exchange reaction design considerations

Preliminary experiments were performed following the halogen exchange reaction originated by^45^. Mass spectral analysis of an early natural abundance test reaction showed that trifluoroacetic acid was formed by reaction of AgBF_4_ with CBr_3_CO_2_H in DCM. However, ^13^C NMR multiplet analysis of a test reaction with ^13^CBr_3_CO_2_H showed a conversion to ^13^CF_3_CO_2_H of only 18% after stirring at for 10 days at room temperature. A longer reaction time in glassware was found to be counterproductive because of failure of containment. AgBF_4_ releases highly aggressive BF_3_ during the reaction. The reaction between BF_3_ and silica gel^46,47^ is useful at the purification stage, however, reaction with ground glass joints may give rise to leakage.

### A procedure for making 2-^13^C-trifluoroacetic acid

The following procedure was designed to avoid the necessity of handling hydrogen fluoride, either as a solvent, reagent, or product. Synopsis: 2-^13^C-ethanol is converted to the tribromoacetaldehyde, 2-^13^C-bromal (hydrate) using a molar excess of bromine, Br_2_. One equivalent of water converts the product mixture to bromal hydrate. Reaction with excess nitric acid at a temperature less than 50 °C converted bromal hydrate to 2-^13^Cbromoacetic acid. This product is isolated and converted to 2-^13^C-trifluoroacetic acid using AgBF_4_ under pressure in dichloromethane. The 2-^13^C-trifluoroacetate is extracted into ammonia. Impure ammonium 2-^13^C-trifluoroacetate can be enhanced in purity by sublimation at 85 °C and with a vacuum less than 10 microns Hg. The yields were very low.

### Conversion of 2-^13^C-ethanol to 2-^13^C-bromal (hydrate)

The reaction proceeds according to:

2-^13^C-ethanol + 4Br_2_ → 2-^13^C-bromal + 5HBr

In a closed system, the above is an equilibrium reaction. To drive the reaction to completion, product HBr gas must be permitted to escape. This loss of mass results in a considerable reduction in the volume of the reaction mixture. We have tried sulfur and I_2_ as catalysts. TFA is probably a better catalyst for this reaction. The oxidation potential of Br_2_ is not sufficient to carry oxidation beyond the aldehyde. The aldehyde is required for tribromination, because each bromination step proceeds via the enol. To avoid loss of volatiles, 2-^13^C-ethanol and excess Br_2_ were combined at liquid nitrogen temperature and warmed very slowly to reflux temperature. When the reaction has been driven to completion, ^13^C NMR shows the presence of only 2-^13^C-bromal (~40 ppm) and its hydrate (~12 ppm). Prior to the next step, it may be desirable to isolate 2-^13^C-bromal via distillation, and its hydrate by crystallization but it is not essential. One equivalent of H_2_O is added to convert all to hydrate.

### Conversion of 2-^13^C-bromal hydrate to 2-^13^C-tribromoacetic acid

^13^CBr_3_CH(OH)_2_ + HNO_3_ → ^13^CBr_3_CO_2_H + ½N_2_O_4_ + H_2_O

The more stable hydrate of tribromoacetaldehyde is the species oxidized. Completion of this reaction can be determined by ^13^C NMR (~34 ppm). At completion of reaction, excess nitric acid is removed under vacuum. Purity can be determined by melting point, 128 °C.

### Conversion of 2-^13^C-tribromoacetic acid to 2-^13^C-trifluoroacetic acid

^13^CBr_3_CO_2_H + 3AgBF_4_ + 3(C_2_H_5_)_2_O → ^13^CF_3_CO_2_H + 3AgBr + 3[BF_3_·(C_2_H_5_)_2_O]

This reaction is conducted in dichloromethane. This three-step reaction is exceedingly slow. For practical synthesis, it is necessary to conduct this reaction in a sealed pressure reaction vessel at above 75 °C. The completion of this reaction can be determined by ^13^C

NMR (quartet at 116.6 ppm is dominant) or ^19^F NMR (doublet at −76.55 ppm is dominant). Under standard conditions BF_3_ is a gas, whereas BF_3_·(C_2_H_5_)_2_O is liquid. A strong Lewis acid, BF_3_ forms a complex with diethyl ether. This reduces the pressure. At the completion of the reaction, the DCM reaction mixture is eluted through silica gel. SiO_2_ reacts with BF_3_. 2-^13^C-trifluoroacetate is extracted into aqueous ammonia and excess removed under vacuum. Ammonium 2-^13^C-trifluoroacetate was converted into a fine powder by lyophilization, to aid in sublimation. Ammonium 2-^13^C-trifluoroacetate was sublimed to improve purity by sublimation at 85°C and less than 10 microns Hg vacuum. Because initial purity was poor, yield was poor. Gram scale quantities of material could be processed in this way.

### Manipulation of ^13^C haloacetates

The halogen exchange reaction between 2-^13^C-tribromoacetic acid and AgBF_4_ proceeds in a stepwise fashion and hence yields a mixture of haloacetates. The target compound, 2-^13^C-trifluoroacetic acid is too volatile and so the haloacetate mixture is best manipulated as a salt. The ammonium salts have volatilities that were exploited for purification by high vacuum sublimation and the progress of purification was monitored by ^19^F and ^13^C NMR. Whereas high vacuum sublimation has only one theoretical plate, the greater load capacity compared to chromatography affords it an advantage for the first stages of purification. Repeated stages of high vacuum sublimation yielded a product that gave three spots on silica gel TLC. Neutral alumina TLC gave streaks. One spot showed an R_f_=0.78 on silica gel TLC that was identical to the R_f_ of natural abundance ammonium trifluoroacetate, eluted with 10% aqueous ammonia in methanol. Based on this observation, the ammonium ^13^C haloacetate mixture was subjected to preparative TLC under the same conditions. ^19^F and ^13^C NMR of the product from the preparative TLC target band showed that the mixture consisted of ammonium salts of 2-^13^Ctrifluoroacetate, 2-^13^C-bromodifluoroacetate and 2-^13^C-dibromofluoroacetate (See Table I).

It is interesting to note that CF_3_I has been enriched to 86% in ^13^C by selective multiphoton dissociation of ^12^CF_3_I at pressures less than 1.0 torr^48^. A more practical method would rely on enrichment in the condensed phase. An isotope effect is often observed on melting points^49^. In recent work, boron has been enriched to 93.21% in ^10^B and 99.01% in ^11^B by zone refining^50^. It is known that isotope effects on heat capacity and crystal transition temperature can be detected by differential scanning calorimetry^51^. Zone refining of low melting ammonium trifluoroacetate (M.P. 123 °C) would be more energy efficient than that of high melting boron (M.P. 2079 °C). In future, we plan to develop an analytical HPLC method and to use preparative HPLC for the purification of ammonium 2-^13^C-trifluoroacetate. Infrared difference spectroscopy may be able to resolve the carbon isotope effect for ammonium 2-^13^C-trifluoroacetate in the condensed phase. Differential scanning calorimetry of ammonium 2-^13^C-trifluoroacetate will provide the thermodynamic information needed to plan and develop a zone refining method for the extraction of ammonium 2-^13^C-trifluoroacetate from an inexpensive natural abundance melt.

Another promising route to ^13^C enriched TFA that we may explore is that of fluorodeoxygenation^52^, starting with glycine. Recently a new and better synthesis of arylsulfur trifluorides has been reported^53^ for reagents that may provide a convenient route to fluorodeoxygenation of carboxylic acids.

## Conclusion

We have presented a critical scientific narrative of a promising technology. The advantage of the ^13^CF_3_ NMR spin label in protein NMR has been explained. Progress in incorporation of this spin label in a perdeuterated amino acid and in an important protein is reported. We comment on some of the synthetic subtleties encountered.

Whereas incorporation of monofluorinated aromatic amino acids in proteins using in vivo or in vitro expression systems is now routine, incorporation of trifluoromethyl analogs of branched chain amino acids is at present a challenging technology. A biochemical hurdle is the specificity of an aminoacyl tRNA synthetase. Wang et al.^24^ determined that the specificity constant, k_cat_/K_M_ of 5,5,5-trifluoroisoleucine is only 1/134 that of isoleucine for E. coli isoleucyl tRNA synthetase. Considerations of enzyme kinetics and competitive inhibition dictate that the background concentration of isoleucine needs to be effectively zero for tRNA^Ile^ to be charged with 5,5,5-trifluoroisoleucine by isoleucyl tRNA synthetase. In addition to the catalytic domain, where the amino acid and tRNA^aa^ are specifically ligated, aminoacyl tRNA synthetase also has an editing domain for the hydrolysis of mischarged tRNA. In the case of isoleucyl tRNA synthetase, the editing domain has been evolved most specifically for the hydrolysis of val· tRNA^Ile^. The successful incorporation of 5,5,5-trifluoroisoleucine in mDHFR, mIL II (24) and in calmodulin^55^ (M. Kainosho: private communication) implies that 5TFI· tRNA^Ile^ is too large for the editing site. Practical questions remain. Is an expression system based upon E. coli more advantageous than one based upon a eukaryotic organism? Is a cell free lysate system preferable to in vivo expression system?

This paper covers many areas of synthetic chemistry, organic, fluoro, isotopic, and biochemical. We have identified ammonium 2-^13^C-trifluoroacetate as an important synthon for introduction of the ^13^CF_3_ group into amino acids. Completed synthesis of methallylcyanide-d_7_ and conceptual synthesis of 5-^13^C-5,5,5-trifluoroisoleucine-d_7_, provide an important building block for the exploitation of ^13^CF_3_ in protein NMR.

5,5,5-TFI is an unnatural AA, more so than monofluorinated amino acids. Either as uniform, or site specific isotopic labels, deuterium and ^13^C amino acids are readily synthesized and incorporated. Fluorine is the 13^th^ most abundant isotope in the earth’s crust, yet even after 3.5b years of biology only about a dozen fluorinated natural products have been evolved, attributed to fluorine’s chemistry as a “superhalogen”^54^. Organofluorine compounds as polymers or as drugs have proven useful in material science and pharmacology. The target spin ^13^CF_3_ label should prove useful in multidimensional heteronuclear NMR structure dynamics studies of proteins. Synthesis of 5,5,5-TFI has been proven by its incorporation in the calcium binding protein calmodulin. Methallylcyanide-d_7_ has been produced with military grade deuterium isotope purity, 99.4%. Trace quantities of 2-^13^C-trifluoroacetate have been characterized by ^13^C-^19^F NMR coupling. This contribution may pave the way to future study.

This work is part of a project in heteronuclear multidimensional NMR^57,58,59^.

**Table 1.**
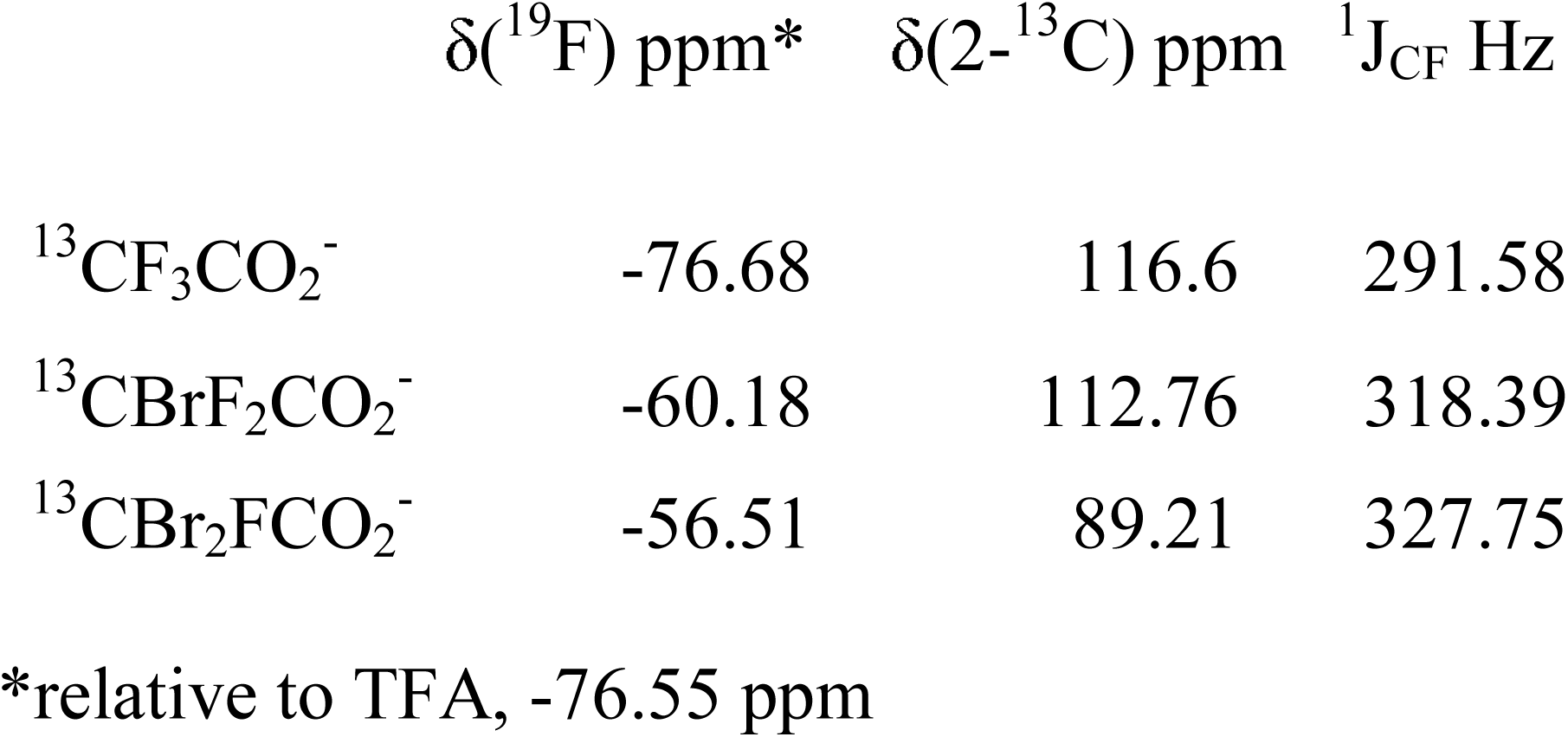

| | $\delta(^{19}\text{F})$ ppm* | $\delta(2\text{-}^{13}\text{C})$ ppm | $^1J_{\text{CF}}$ Hz |
| --- | --- | --- | --- |
| $^{13}\text{CF}_3\text{CO}_2^-$ | -76.68 | 116.6 | 291.58 |
| $^{13}\text{CBrF}_2\text{CO}_2^-$ | -60.18 | 112.76 | 318.39 |
| $^{13}\text{CBr}_2\text{FCO}_2^-$ | -56.51 | 89.21 | 327.75 |

\*relative to TFA, -76.55 ppm

